# Caffeine boosts preparatory attention for reward-related stimulus information

**DOI:** 10.1101/697177

**Authors:** Berry van den Berg, Marlon de Jong, Marty G. Woldorff, Monicque M. Lorist

## Abstract

Both the intake of caffeine-containing substances and the prospect of reward for performing a cognitive task have been associated with improved behavioral performance. To investigate the possible common and interactive influences of caffeine and reward-prospect on preparatory attention, we tested 24 participants during a 2-session experiment in which they performed a cued-reward color-word Stroop task. On each trial, participants were presented with a cue to inform them whether they had to prepare for presentation of a Stroop stimulus and whether they could receive a reward if they performed well on that trial. Prior to each session, participants received either coffee with caffeine (3 mg/kg bodyweight) or with placebo (3 mg/kg bodyweight lactose). In addition to behavioral measures, electroencephalography (EEG) measures of electrical brain activity were recorded. Results showed that both the intake of caffeine and the prospect of reward improved speed and accuracy, with the effects of caffeine and reward-prospect being additive on performance. Neurally, reward-prospect resulted in an enlarged contingent negative variation (CNV) and reduced posterior alpha power (indicating increased cortical activity), both hallmark neural markers for preparatory attention. Moreover, the CNV enhancement for reward-prospect trials was considerably more pronounced in the caffeine condition as compared to the placebo condition. These results thus suggest that caffeine intake boosts preparatory attention for task-relevant information, especially when performance on that task can lead to reward.

---

Humans have always searched for ways to enhance their cognitive capacities in order to efficiently cope with the overload of information they encounter in everyday life. One of the most useful set of mechanisms humans have available to distinguish between relevant and irrelevant information is our attention system (Petersen & Posner, 2012). In particular, this system selectively guides attention towards environmental stimuli and events that are associated with the possibility of gaining reward (Aarts, van Holstein, & Cools, 2011; Engelmann & Pessoa, 2007; Hickey, Chelazzi, & Theeuwes, 2010; Hickey & van Zoest, 2012; van den Berg, Geib, Martín, & Woldorff, 2019; Watanabe, Roelfsema, & van Ooyen, 2010) In addition, caffeine-containing substances (e.g., coffee, tea), arguably the most widely used psychoactive cognitive enhancer in the world, are being used on a daily basis to boost this attention system (Lorist & Tops, 2003; Saville, de Morree, Dundon, Marcora, & Klein, 2018; Wilhelmus et al., 2017).

Both the prospect of gaining reward and the consumption of caffeine-containing substances have been found to improve behavioral performance. For instance, cueing of the prospect of gaining monetary rewards has been found to guide attention *selectively* towards the potentially rewarding events or stimuli (Schevernels, Krebs, Santens, Woldorff, & Boehler, 2014; van den Berg, Krebs, Lorist, & Woldorff, 2014). Neurally, the prospect of gaining reward on an upcoming visual task triggers increased activity in cortical regions involved in attentional control and to improved processing of relevant information in the visual cortices on that task, thereby improving behavioral performance (Hickey et al., 2010; van den Berg et al., 2019; Watanabe et al., 2010). Similarly, caffeine, compared to placebo, has also been found to increase neural activation in cortical areas involved in the selection of relevant information. Specifically, caffeine biases the processing of relevant stimuli over irrelevant ones, including enabling more effective suppression of irrelevant information (Lorist et al., 1995, 1995, 1996; Ruijter et al., 2000a). In addition to these task specific effects, several studies have also showed a broad sustained increase in neural arousal, after the intake of caffeine (vs. placebo), consistent with the more general stimulating effects of caffeine on behavior (Kenemans and Lorist, 1995). As a result of these neural modulations, in habitual coffee drinkers caffeine doses as low as the dose in half a cup of coffee have been found to speed up reaction times (RTs) and improve accuracy (Lieberman, Wurtman, Emde, Roberts, & Coviella, 1987).

Although both reward-prospect and caffeine intake have substantial beneficial effects on behavioral performance, the nature of their interaction has remained elusive. There is substantial evidence, however, that both reward-prospect and caffeine influence behavior through an effect on attentional preparation. In studies employing high spatial resolution fMRI and ones using high temporal resolution EEG recordings, attentional preparation has been linked to activity in the fronto-parietal control regions (reviewed by Corbetta and Shullman 2000). These brain regions are thought to be the main contributors of the fronto-centrally distributed contingent negative variation (CNV)(Grent-’t-Jong & Woldorff, 2007; Walter, Cooper, Aldridge, McCallum, & Winter, 1964), a negative-polarity slow-wave event-related potential (ERP) that is elicited before an imperative stimulus is presented (Brunia, van Boxtel, & Bocker, 2012), and has been found to be a good index of preparatory attention. Another neural marker indexing preparatory attentional processes is oscillatory activity in the alpha-band frequency range (8 – 14Hz). Decreases of the power of this frequency band have been related to increased cortical activity and enhanced selective attention (Scheeringa, Petersson, Kleinschmidt, Jensen, & Bastiaansen, 2012; Worden, Foxe, Wang, & Simpson, 2000). In addition, both the CNV amplitude and the power in the alpha frequency band have been shown to predict behavioral performance (Hillyard, 1969; van den Berg, Appelbaum, Clark, Lorist, & Woldorff, 2016), and they have previously both separately been found to be modulated by reward-prospect and by caffeine (Ashton, Millman, Telford, & Thompson, 1974; Kenemans & Lorist, 1995; Tieges, Snel, Kok, Plat, & Ridderinkhof, 2007; van den Berg et al., 2014).

The aim of the present study was to examine whether and how the prospect of reward and the consumption of caffeine-containing substances, two factors that both enhance attention in everyday life, interact on both the behavioral and neural levels. To investigate this interaction, participants participated in a two-session study in which they either received coffee with caffeine or with lactose (placebo) prior to the experimental session, in which they performed an adapted version of the cued-reward task of van den Berg and colleagues (2014). In this task (**Figure 1**), participants were instructed to respond as fast and accurately as possible to target Stroop stimuli. At the beginning of each trial, participants were presented with a cue that indicated that there was prospect of receiving a reward on that trial or there was no such prospect. During reward-prospect trials participants could earn money if they responded accurately and sufficiently fast.

**Figure 1:**
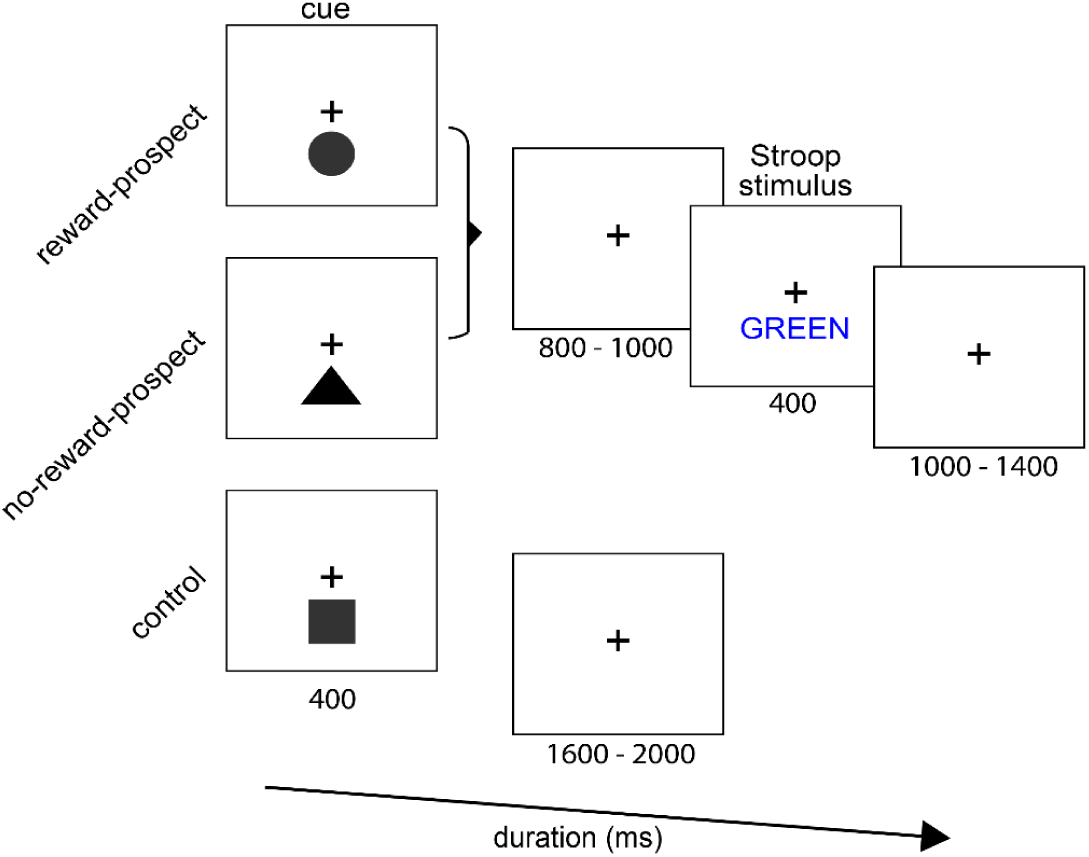
Task and stimuli.

Based on previous findings, our first main hypothesis was that both reward-prospect and caffeine would improve behavioral performance. Neurally, we hypothesized they would both enhance preparatory cortical activity, as indexed by the CNV and alpha power. In addition, we had two main competing hypothesis in terms of potential interactions of these factors. On the one hand, previous research has indicated that caffeine specifically improves selective attention towards relevant information (Lorist, Snel, Kok, & Mulder, 1994; Lorist, Snel, Mulder, & Kok, 1995; Ruijter, Lorist, Snel, & De Ruiter, 2000). Based on these studies, one might expect that caffeine would lead to enhanced attentional preparation for more important stimuli (i.e., in reward-prospect trials) as compared to less important ones (i.e., in no-reward-prospect trials). On the other hand, there is evidence that the stimulating effects of caffeine are most pronounced in situations when attentional control of perceptual functions is reduced, such as in the presence of mental fatigue or a lack of motivation (Koelega, 1993; Lorist et al., 1994; Ruijter, Lorist, & Snel, 1999; Weiss & Laties, 1962). Based on these findings, an alternative hypothesis is that the effects of caffeine would be most pronounced in the no-reward-prospect condition compared to the reward-prospect one, where the attentional system is already triggered by the anticipation of reward. Finally, we inspected the effect of the factors caffeine and reward on the processing of the Stroop stimulus as indicated by the late positive complex (LPC), a component that indicates the processing of target stimulus information (Kappenman & Luck, 2012; van den Berg et al., 2014). Because the LPC has generally been found to be related to response speed, for the LPC we expected the amplitude to parallel the effects of reward and caffeine on behavioral performance.

## Results

### Behavioral performance

Participants responded on average 58 ms (se = 5 ms) more slowly and 2.9 % (se = 0.5 %) less accurately to incongruent than to congruent Stroop stimuli, replicating a multitude of studies of the behavioral effects of Stroop incongruency (MacLeod, 1991) (main effect of congruency; RT: F(1,25) = 203.66, p < .001; accuracy: χ2(1) = 77.79, p < .001). In addition, they responded more quickly and more accurately to the font-color of the Stroop stimulus if the cue for that trial indicated reward-prospect as compared to the no-reward-prospect (main effect of reward; RT: F(1,25) = 44.83, p < .001; accuracy: χ2(1)=13.34, p < .001; **Figure 2**). Moreover, we found that RTs decreased and the proportion of hits increased in the caffeine condition compared to the placebo condition (main effect caffeine; RT: F(1,40802) = 232.21, p < .001; accuracy: χ2(1) = 139.47, p < .001; **Figure 2**). On the behavioral level, no significant interactions were observed between the independent variables of congruency, reward, and caffeine.

**Figure 2:**
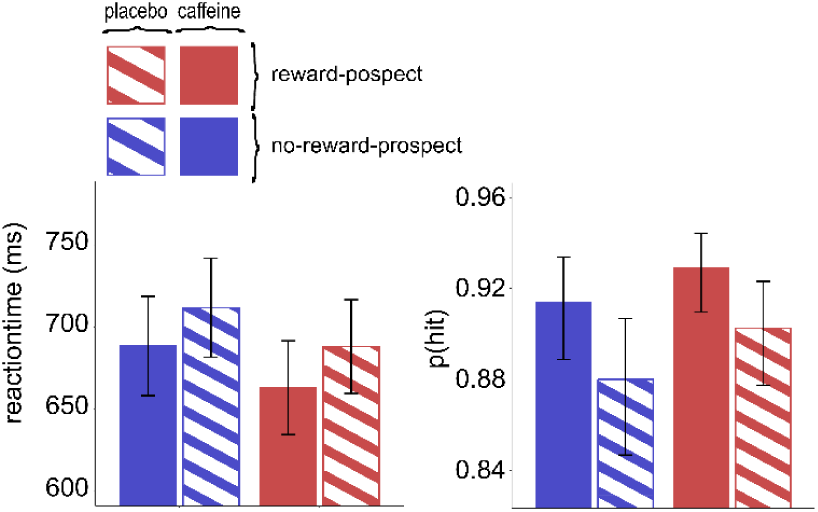
Behavioral results. Participants responded faster and more accurately when cued with reward-prospect vs. no-reward-prospect and when receiving caffeine vs. placebo. Error bars reflect 95 % CIs.

### EEG results

#### Cue-evoked brain activation: Influence of reward and caffeine on the attention-sensitive CNV

Visual inspection of Figure 3 shows that the ERPs evoked by both the reward-prospect and no-reward-prospect cues started to diverge from the control trials around 700ms after cue onset. In the conditions for which the participants were cued that an upcoming Stroop stimulus would appear, a more pronounced fronto-central slow-wave negative deflection (the CNV) was observed compared to the control condition (in which the participant knew no Stroop stimulus would be coming). This CNV was more negative following a reward-prospect cue than to the no-reward-prospect cue, replicating the results of van den Berg et al., (2014) (main effect of reward: F(1,25) = 9.84, p = .004). Whereas we did not observe a significant interaction between reward and caffeine on the behavioral measures, the effects of caffeine on CNV amplitude were dependent on the reward condition. The CNV following a reward-prospect cue was larger when participants had received caffeine prior to the experiment as compared to placebo, while this CNV difference between caffeine and placebo was absent in the no-reward-prospect trials (**Figure 3 A and B**) (reward x caffeine interaction: F(1,45871) = 8.31, p = .004; reward-prospect ^*caffeine* **minus** *placebo*^: t(45862) = −4.31, p < .001; no-reward-prospect ^*caffeine* **minus** *placebo*^: t(45971) = -.23, n.s.).

**Figure 3:**
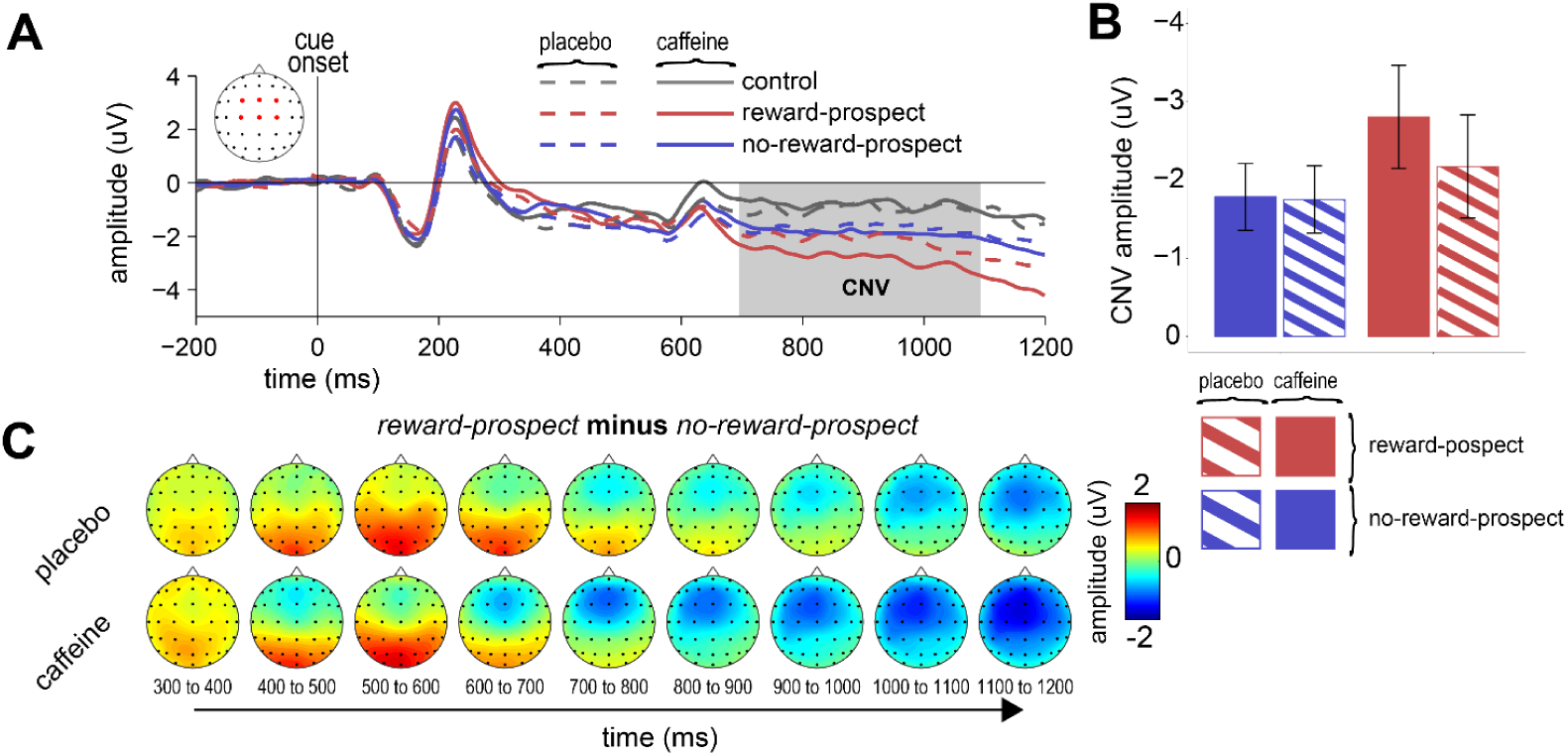
Cue-evoked ERPs**. A.** Grand average cue-evoked responses from fronto-central ROIs for the different conditions. **B.** CNV amplitude for the different conditions. **C.** Differences in the topographical distributions of the no-reward rewardprospect versus control and reward-prospect versus no-reward-prospect conditions.

#### Cue-evoked brain activation: Influence of reward and caffeine on alpha power

The frequency analyses revealed lower levels of sustained alpha power (8 to 14 Hz) (i.e., sustained across the whole session) over occipital brain regions if participants received coffee with caffeine prior to the experiment as opposed to coffee with placebo (**Figure 4A**) (main effect of caffeine: F(1,57260) = 1803.59, p < .001), replicating previous work (Kenemans & Lorist, 1995). In addition to these sustained effects of caffeine, we observed cue-evoked changes in alpha power over occipital electrodes. More specifically, after presentation of the cue, occipital alpha power decreased more in the caffeine condition (reflecting increased preparatory attention) as compared to the placebo condition (main effect of caffeine, ROI^o^: F(1, 45848) = 992.95, p < .001; **Figure 4B and C**). In addition, alpha power decreased in response to reward-prospect cues compared to no-reward-prospect ones (main effect of reward: F(1, 25) = 12.93, p = .001; **Figure 4B and C**), replicating previous results for this preparatory-attention effect (van den Berg et al., 2014). In contrast to the CNV effects, no interactions between the effect of caffeine and reward was observed in terms of cue triggered alpha power (F(1,45871) = 0.01, n.s.).

**Figure 4:**
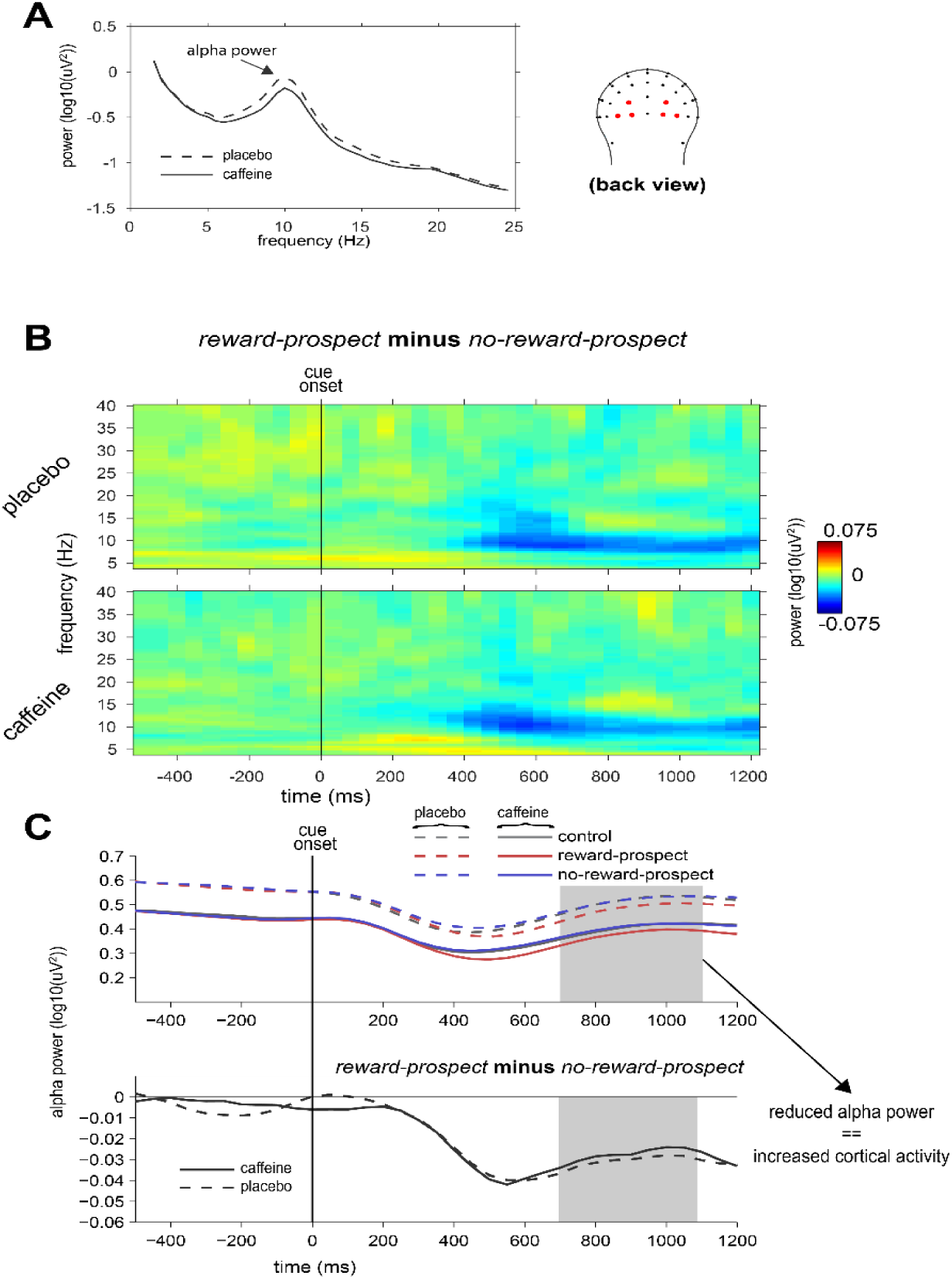
Sustained and cue-locked changes in oscillatory power measured over the occipital channels. **A**. Sustained EEG power across the entire session revealed lower alpha power when participants received caffeine vs. placebo. **B**. Cue-locked spectral power revealed a decrease in alpha power prior to the presentation of the imperative Stroop stimulus, irrespective of caffeine condition. **C.** Changes in alpha power over time relative to the onset of the cue-stimulus.

#### Target-stimulus-evoked brain activation: Influence of reward and caffeine on the LPC ERP component

We observed more pronounced positive ERP amplitudes in both the caffeine and reward-prospect conditions starting around 300 ms at the fronto-central and parietal ROIs compared to the placebo and no-reward-prospect trials, respectively (ROI_fc_: 300 – 500 ms; main effect of caffeine: F(1, 40833) = 331.87, p < .001; main effect of reward: F(1,25) = 29.08, p < .001; ROI_p_: 300 – 500 ms; main effect of caffeine: F(1, 40826) = 197.80, p < .001; main effect of reward: F(1,25) = 8.80, p < .007). Visual inspection of Figure 5 shows that the largest positive-polarity activity was observed in the condition in which the participant both received caffeine and was cued with reward-prospect, while the lowest positivity was elicited in the no-reward-prospect condition during which the participant received placebo, an effect that was confirmed by the statistical analyses in both the fronto-central and parietal ROI (caffeine × reward interaction in the P3 latency range; ROI_fc_: (F(1,40833) = 4.16, p = .041); ROI_p_: (F(1,40833) = 6.94, p = .008). In the fronto-central ROI we observed an effect of reward in both the caffeine condition (caffeine _*reward-prospect* **minus** *no-reward-prospect*_: t(32) = 5.79, p < .001), and a significant effect of reward-prospect in the placebo condition (placebo _*reward-prospect* **minus** *no-reward-prospect*_: t(33) = 4.32, p < .001).In the parietal ROI we observed an effect of reward in the caffeine condition (caffeine _*reward-prospect* **minus** *no-reward-prospect*_: t(37) = 3.77, p < .001), while there was no observable effect of reward-prospect in the placebo condition (placebo _*reward-prospect* **minus** *no-reward-prospect*_: t(38.) = 1.62, n.s.).

**Figure 5:**
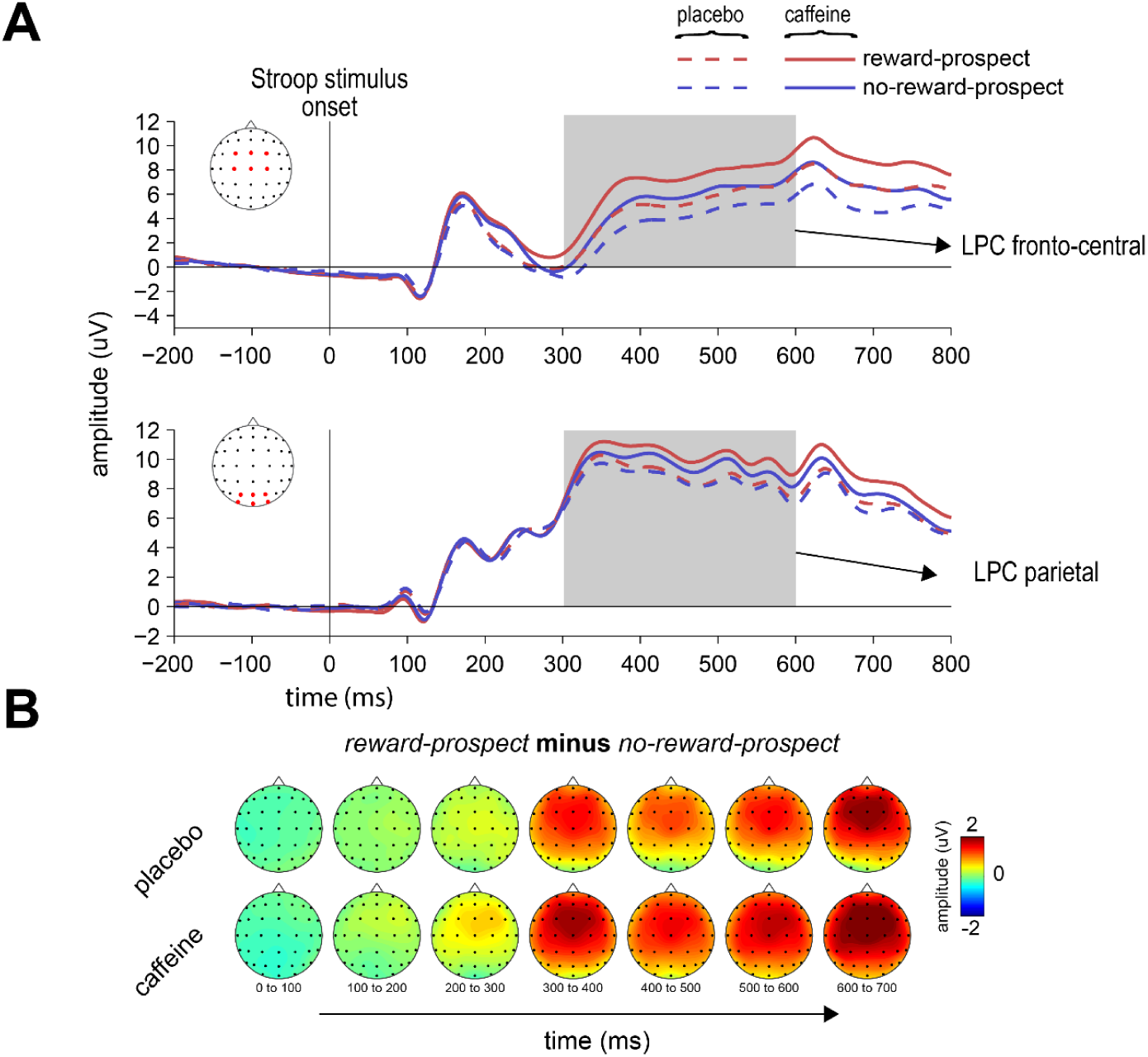
Stimulus-evoked ERPs (collapsed over congruency) for which the participant gave the correct response. **A.** ERP traces from the fronto-central (top panel) and parietal (lower panel) ROIs, showing the large late positive complex (LPC) wave. **B.** Topographical distribution of the difference in ERP amplitude between the reward-prospect and no-reward-prospect trials for the caffeine and the placebo conditions separately, showing the effects of these manipulations on the LPC.

## Discussion

Two factors that are separately known to influence preparatory attention are caffeine and the prospect of reward. The goal of the present study was to investigate if and how these factors interact to enhance attentional preparatory activity and subsequently improve the processing of and response to task-relevant information that can potentially leads to monetary rewards. To do this, we conducted a two-session experiment in which participants performed a cued-reward Stroop task, while behavioral and neural-activity measures were acquired. Each trial of the task consisted of a cue that indicated whether a color-word Stroop stimulus would or would not follow, and whether there was a prospect of gaining monetary rewards for good performance on discriminating that color-word Stroop stimulus. Before the start of each experimental session, participants received either a cup of decaffeinated coffee with caffeine or with lactose (placebo) (both 3 mg/kg bodyweight). The key results showed that caffeine intake indeed resulted in greater enhancements of cue-triggered attention-related neural processes and better behavioral performance, especially when the cue indicated that there was a potential for reward for good performance on the subsequent Stroop task. The findings described in this paper advance our understanding of how two previously separately studied factors, both of which impact behavioral performance, interact on a neural level to improve behavioral performance even further.

### General effects of caffeine – increased neural activity and improvements in behavioral performance

The level of general neural arousal substantially increased after the intake of caffeine (vs. placebo). Replicating previous studies, we observed reductions of power in the alpha frequency band (8 to 14 Hz), reflecting an increase in cortical brain activity and alertness (Kenemans & Lorist, 1995; Scheeringa et al., 2012). This increased state of alertness when participants had received caffeine prior to the start of the experimental session, was paired with improved behavioral performance (both RTs and accuracy), suggesting that caffeine intake improves the general state of the brain to such an extent that individuals can react more effectively to external stimuli and events.

### Behavioral improvements by both caffeine and reward

In line with previous research, the behavioral responses to the color-word Stroop stimuli were faster and more accurate when preceded by a cue that indicated a potential for reward for that trial (Padmala & Pessoa, 2011; Schevernels et al., 2014; van den Berg et al., 2014). The effects of reward-prospect and caffeine appeared to be additive in terms of RTs and accuracy, and thus behavioral performance was most optimal when there was both a prospect of reward and caffeine had been ingested. In contrast to these results, some previous studies have reported that caffeine is especially effective when, due to the subject’s state, behavioral performance was *<u>not</u>* optimal. For instance, it has been found that caffeine can have a profound impact on response times and accuracy particularly when participants are fatigued (Lorist & Tops, 2003; Nehlig, 2010). Mental fatigue is a subject state that has been associated with decreased behavioral performance, usually after continuously performing a taxing task for an extended time period (Lorist and Faber, 2013). Thus, our finding that caffeine actually boosted behavioral performance similarly in both the more optimal reward-prospect condition and the suboptimal no-reward-prospect condition would seem to differ from previous reports.

### Improvements in attentional-preparation and target-processing by caffeine and reward

Within session, on a trial-by-trial basis, increased behavioral performance was preceded by enhancements in processes related to event-related attentional preparation. These effects were modulated by caffeine and reward, resulting in enhancements of both the fronto-central CNV and posterior alpha power starting at ∼700 ms following the cue, reflecting the marshalling of neural circuits that have been associated with preparatory attention (Corbetta & Shulman, 2002; Hillyard, 1969) and the enhancement of sensitivity to visual information (i.e., alpha power) (van den Berg et al., 2019; Worden et al., 2000), after caffeine consumption and during reward-prospect trials.

From a theoretical perspective, reward-prospect has been thought to enhance the saliency of specific impending events (Schevernels et al., 2014; van den Berg et al., 2014), resulting in the recruitment of the attentional-control circuits to improve the processing of those events. Similar to the improvements in preparatory attention by the reward-prospect cue and by caffeine, the processing of the impending Stroop stimulus was also improved by both of these factors. The effect of cueing with the potential for obtaining a monetary reward (vs. no-reward) manifested as a larger fronto-central distributed positivity (LPC) starting at 300 ms, similar to the effect observed by van den Berg et al., (2014). Again, this enhancement was larger when caffeine was administered prior to the start of the experiment. The increase in size of the LPC by caffeine and reward-prospect could reflect increased quality of processing of the Stroop stimulus (Kappenman & Luck, 2012; van den Berg et al., 2014).

### Differential neural markers of attentional preparation for reward-prospect and caffeine

In the present study, we used the CNV amplitude and occipital alpha power as markers for cue-triggered preparatory attention. First, we replicated that both caffeine and reward-prospect have an enhancing effect on the slow-wave CNV (Schevernels et al., 2014; van den Berg et al., 2014) and on posterior alpha power (van den Berg et al., 2014). Second, and probably even more intriguingly, we observed that the modulation of the CNV by reward-prospect could be increased even further after the intake of caffeine. It is important to note that in previous studies, the CNV amplitude and alpha power have been statistically linked through correlation to improved target processing and more optimal behavioral performance (Grent-’t-Jong, Boehler, Kenemans, & Woldorff, 2011; van den Berg et al., 2016, 2014). Hence, one might expect that the CNV and alpha power could be reflective of the same underlying neural mechanism that is involved in attentional preparation. However, this hypothesis did not seem to hold in these studies, since no correlation between the CNV amplitude and alpha power was observed on a trial-by-trial basis.

Here, we found that the effects of reward and caffeine interacted on the CNV but were additive for alpha power. This dissociation suggests that these markers likely reflect windows into two different facets of preparatory attention. Although their specific neural and cognitive functions are not fully understood, the CNV has been suggested to reflect task focused attentional preparatory processes originating from the fronto-parietal attentional control circuits (Grent-’t-Jong & Woldorff, 2007). Decreases in occipital alpha power have been correlated with increased BOLD signal in the visual cortices (Scheeringa et al., 2012), suggesting that the amplitude of this pre-target oscillatory activity is inversely reflective of receptibility to information in the visual sensory cortices. In the present study, we found that, on the one hand, caffeine influenced the neural circuitry underlying the CNV when cued with reward-prospect but not when cued with no-reward-prospect. On the other hand, alpha power was influenced by caffeine regardless of cue information. These results suggest that caffeine may have a more direct impact on attentional preparatory processes for upcoming relevant stimuli than previously thought. Continuing the line of reasoning that caffeine can enhance the selectivity of processing of specific events, these findings suggest that caffeine works on our neural information processing system in at least two ways. First, by enhancing the general state of the subject, caffeine increases processing capabilities (perhaps as reflected by lower sustained alpha power). Second, the effect of caffeine depends on the context of an event, that is, on its behavioral relevance. Neurally this can be illustrated by a larger CNV to reward-predicting cues under the influence of caffeine (vs. placebo), while there is little effect of caffeine on the CNV triggered by cues that predict no-reward. Our finding that the prospect of gaining reward and the consumption of caffeine speeded up reactions to subsequent Stroop stimuli in an additive manner confirms the differential effect these factors on our attention system at the underlying neural level.

Previous caffeine studies have found a pronounced effect of caffeine when mental fatigue occurs (and thus when slower RTs would have been observed), indicating that caffeine enhances the state of the subjects in such a way that it helps them overcome (or at least compensate for) low levels of neural arousal that tend to impair behavioral task performance. In line with these observations, we found that caffeine did indeed seem to improve the general arousal state of the subject (as indexed by lower alpha power, thus reflecting higher levels of cortical activation). Here, we manipulated the behavioral importance of events through reward-prospect cues. Under these circumstances we found that caffeine resulted in greater enhancement of attention for the more important events. Thus, in addition to the more general effect, caffeine can specifically boost attentional preparation for more salient or behaviorally important external events (as signaled by the reward-prospect cue) compared to other events that are less important. These findings suggest that improved behavioral performance due to the enhanced state induced by caffeine might depend on the context, and that this context is behavioral relevance.

In addition to these effects of caffeine and reward, our behavioral findings showed that participants performed faster and more accurately after presentation of congruent as compared to incongruent Stroop stimuli, consistent with a multitude of previous studies using the Stroop task (MacLeod, 1991). This congruency effect was not modulated by reward-prospect when averaged across subjects, replicating van den Berg et al (2014). We should note that in the present study our main question was not focused on the interaction between reward-prospect and stimulus-conflict processing. However, given the mixed results of previous studies that have looked at this question (see for instance van den Berg et al., 2014), future replication studies should be conducted to further investigate the nature of the relationship between reward-prospect and stimulus-conflict processing. Furthermore, we did not find an interaction between caffeine and congruency, similar to previous research (Kenemans, Wieleman, Zeegers, & Verbaten, 1999; Tieges et al., 2007)

### Conclusion

In the present study, we found that reward-prospect and caffeine intake can enhance neural attentional-preparatory activity, showing most optimal preparation and behavioral performance after caffeine intake and after a reward-prospect cue, as reflected by larger CNV and lowest alpha power being triggered by the cue. Additionally, we found that caffeine appears to especially improve preparatory attention for a task that could potentially result in a reward In a broader sense these findings indicate that caffeine can specifically target attentional preparatory neural processes for important salient events relative to events that are less consequential. These improvements in preparatory processes then result in better target processing and more optimal behavioral performance.

## Materials and Methods

### Participants

Thirty-one healthy adults (10 males), ranging in age from 18 to 31 (M = 22.0 year, SD = 3.6), participated in the two-session experiment. Participants received either course credits or 7 euros per hour for participation. In addition, they received a monetary reward that depended on their performance (M = 10.6 euros, SD = 3.9). All participants were native Dutch speakers, right-handed, regular coffee-drinkers who ingested a minimum of 2 cups per day (M = 4.2 cups/day, SD = 1.7). They had a regular sleep schedule and had normal or corrected-to-normal-visual acuity. Participants indicated that they were not lactose-intolerant and that they did not smoke. Data from two participants was excluded due to technical issues during recording, while data from another three were excluded due to excessive noise in the EEG (i.e., > 30% of the EEG epochs rejected due to artifacts, see EEG preprocessing below). The experiment was approved by the Ethics Committee of the Psychology Department of the University of Groningen, and participants gave their written informed consent before the start of the first experimental session.

### Apparatus

The experiment was conducted in a sound- and light-attenuated room with the stimuli being presented on a 100Hz LCD monitor with a resolution of 1920 × 1080 (Iiyama ProLite G2773HS). Participants sat in a comfortable desk chair at a viewing distance of 70 cm from the monitor and gave behavioral responses using a gamepad with four bumper buttons, using the index and middle finger of their left and right hands (Logitech Rumblepad, http://www.logitech.com/). The experimental task was programmed using the Presentation software package (version 18.1 06.09.15, http://www.neurobs.com/). Stimuli were randomized using the R statistical programming software package (R Development Core Team, 2013).

### Task and Stimuli

During the entire task, a central fixation cross was continually visible in the middle of the screen. At the start of each trial (**Figure 1**), a cue stimulus (circle, triangle, or square [visual angle: 1.23°]) was presented, 1 cm below the fixation cross, for 400 ms. This cue stimulus indicated whether the trial was a reward-prospect trial (40 % of the trials), in which a monetary reward would be given if a response to the subsequent imperative stimulus (a Stroop stimulus) was both correct and met a pre-defined response time (RT) criterion (see below), or whether was a no-reward-prospect trial (40 %) or a control trial (20 %), in which no imperative Stroop stimulus followed the cue. The meaning-shape mapping of the cue stimuli was counterbalanced across participants. Following a fixation screen presented for 800 to 1000 ms, in the reward-prospect and no-reward-prospect trials, a Stroop color-word stimulus (i.e., Dutch words for “RED”, “GREEN”, “BLUE”, and “YELLOW” [visual angle: 1.23° by 4.91°]) was presented below the fixation mark for 400 ms. Participants were instructed to indicate the font-color of the Stroop stimulus fast and accurate. In half of the trials, the Stroop stimulus was congruent (i.e,., word meaning matched the font color) and in the other half the stimuli were incongruent (i.e., word meaning did not match the font color). The interval between the offset of the Stroop stimulus and the next cue varied randomly between 1000 and 1400 ms. The interval between the control cue (i.e., the one indicating that no Stroop target stimulus would follow) and the cue for the next trial was varied randomly between 1600 and 2000 ms.

Participants were instructed to respond to the font-color of the Stroop stimulus as fast and as accurately as possible, by pressing one of the four buttons corresponding to the font-color on the gamepad. After receiving instructions, the participants first performed a practice block of 30 trials, in which they received positive feedback (‘correct’) if their response was correct and faster than 900 ms, or negative feedback (‘incorrect’) if the response was incorrect or slower than 900 ms. If participants did not achieve a hit-rate of over 80 %, they performed a second practice block of 30 trials; otherwise the experimental task started.

After the practice trials, participants performed a block of 30 trials, which was used to calculate the RT criterion (RT_crit =_ mean RT + 200 ms) in the reward-prospect condition. The number of points participants could earn was based on the RT on each individual trial (RT_i_ in ms) according to the formula: RT_crit_ - RT_i_. These points were converted to euros (i.e., 3000 points represented 1 euro). Participants did not receive a penalty if they responded too slowly or incorrectly, and hence they could only gain money and not lose any.

Thereafter subjects performed 6 experimental blocks of 200 trials each. After each block participants could to take a self-timed break. Within each block there were 5 s breaks every 15 trials and a 30 s break every 100 trials. After every 30 trials a screen was presented for 2000 ms. that provided feedback, namely indicating the total amount of money made thus far.

### Procedure

The experiment consisted of two sessions that were scheduled exactly one week apart. Both sessions started at 9:00 a.m. and each took approximately 3.5 hours. Participants were instructed to abstain from alcohol and caffeine-containing substances for at least 12 h before each session. Approximately 45 min before the start of the task, participants received a cup of decaffeinated coffee, to which either caffeine or lactose (both 3 mg/kg bodyweight [bodyweight was reported by the participant]) had been added. The experimenter was blind to whether caffeine or lactose had been added. In addition, participants were not informed that the coffee could contain either caffeine or lactose. The order of both conditions was counterbalanced over sessions across participants.

### EEG recording and data analysis

EEG was recorded using a 64-channel ANT waveguard electrode cap (10-10 system), using an online average reference. The sampling rate was 512 Hz and the data was filtered during recording using a FIR filter with a corner frequency at 102 Hz (0.2 x sampling rate). Vertical and horizontal EOG activity was recorded from two electrodes placed above and below the right eye and from two electrodes placed lateral to the outer canthi of the two eyes, respectively. Electrode impedances were kept below 5 kΩ. The analyses were performed using custom Matlab scripts (MATLAB - Release 2015b) in combination with the EEG analysis toolboxes Fieldtrip (Oostenveld, Fries, Maris, and Schoffelen, 2011) and EEGlab (Delorme and Makeig, 2004).

Data was offline re-referenced to the algebraic average of the mastoid electrodes. Channels that contained excessive noise were replaced by interpolated values of the surrounding electrodes (spherical spline interpolation). Eye blinks and eye movements were corrected using independent component analysis (ICA) to reconstruct the data excluding those components that reflected eye blinks. The data was filtered using a 0.01 Hz high pass filter. Epochs were extracted from −1500 to 2500 ms surrounding the onset of the cue stimulus and from −1500 to 2500 ms surrounding the onset of the Stroop stimulus. Epochs containing any remaining artifacts (amplitude > 150 µV, −500 to 1500 ms surrounding cue and stimulus onset) were excluded from the analysis (average epochs rejected per subject per session, cue epochs rejected: M = 7.2 % (SD = 7.6 %); target epochs rejected: M = 5.9 %, (SD = 6.3 %)).

After artifact rejection, the mean number of epochs per condition for the cue was as followed: on average each session consisted of 455 (SD = 41) reward-prospect trials, 453 (SD = 39) no-reward-prospect trials, and 219 (SD = 21) control trials. The mean number of epochs per condition for the target was as followed: on average each session consisted of 461 (SD = 39) reward-prospect trials, and 458 (SD = 37) no-reward-prospect trials.

### Event Related Potentials

To statistically examine cue-evoked brain activation, the mean amplitude of the CNV was derived from a fronto-central region of interest (ROI_fc_: FCz, FC1, FC2, Cz, C1 and C2), consistent with previous literature (Brunia et al., 2012), measured in the 700 – 1100 ms interval after cue presentation for every trial. Stimulus-evoked brain activation was examined in two ROIs, namely the mean amplitude of the LPC component (van den Berg et al., 2014) was derived from the fronto-central ROI in the 300 – 600 ms post-stimulus interval, and, in the same interval, the LPC component was extracted from a parietal ROI (ROI_p_: Pz, P3, P4, POz, PO3 and PO4).

### Power Analysis

To investigate sustained changes in the power at different frequencies, fast Fourier transformations (FFT) were performed on the epoched data, using a Hanning window as implemented in Fieldtrip (Oostenveld *et al.*, 2011). To obtain time-locked changes in power, time-frequency decomposition was done by multiplying the epoched data with a moving Hanning window and analyzing the product using a Fourier transform as implemented in Fieldtrip (Oostenveld *et al.*, 2011). Log_10_ transformed power spectra were subsequently binned for each subject and averaged according to the various conditions. No baseline correction was performed for the oscillatory power analyses. To assess the effects of reward and caffeine on alpha activation, we measured alpha power (converted to log10 [uV^2^]) in an occipital ROI (ROI_o_: PO7, PO8, PO3, PO4, O1, O2), consistent with previous studies of attention-related alpha activity (Jensen & Mazaheri, 2010; van den Berg et al., 2014; Worden et al., 2000) between 700 and 1100 ms after cue presentation.

### Statistical analysis

For statistical significance testing, we employed a mixed-modelling approach using the *lme4* statistical package (Bates *et al.*, 2015). The data provided to the model included the response time, and a mean amplitude of the various neural measurements on each trial (i.e., the data consisted of ∼62400 observations [2 sessions × 1200 trials × 26 subjects]). A mixed modelling approach has the advantage of taking into consideration the individual data points (as opposed to averaging the various conditions per subject) and the ability to account for variance that is related to between session effects (for instance, learning effects). The models contained a varying intercept per subject and per session. In addition, a varying slope for reward-prospect by subject was added to the model if the fit of the model improved as indicated by the Akaike Information Criterion (AIC: estimate of the quality of a model relative to other models). To obtain information about statistical significance, the degrees of freedom were approximated using the Satterthwaite approximation of degrees of freedom as calculated by the R package *lmertest* (Kuznetsova *et al.*, 2017). Accuracy was modeled using a logistic regression), and statistical information was obtained using the likelihood ratio test by estimating two models (one with and one without the factor of interest). For visualization purposes logit values for each condition were converted to probabilities (logit_-1_). Statistical tests were considered significant at p < .05. The formulas in Table1 reflect the models used to statistically test the effects of congruency, caffeine, and reward on the behavioral and neural dependent variables.

**Table1:** Models used to statistically test the effects of congruency, caffeine, and reward-prospect on the behavioral and neural dependent variables (as indicated by [variable]_n_ where n reflects trial n). The notation for these models allowed for a varying intercept and a varying slope for each participant (as indicated by j).

|  |  |
| --- | --- |
| Behavioural Analysis | $p(\text{hits})_n = \text{logit}^{-1}(\beta_{j[n]}^0 + \beta^1 \cdot \text{congruency}_n \times \beta^2 \cdot \text{reward}_{j[n]} \times \beta^3 \cdot \text{caffeine}_n + \beta^4 \cdot \text{session}_n + \epsilon_n)$ |
| | $RTs_n = \beta_{j[n]}^0 + \beta^1 \cdot \text{congruency}_{j[n]} \times \beta^2 \cdot \text{reward}_{j[n]} \times \beta^3 \cdot \text{caffeine}_n + \beta^4 \cdot \text{session}_n + \epsilon_n$ |
| ERP analysis | $ERPs(CNV, LPC)_n = \beta_{j[n]}^0 + \beta^1 \cdot \text{reward}_{j[n]} \times \beta^2 \cdot \text{caffeine}_n + \beta^3 \cdot \text{session}_n + \epsilon_n$ |
| Power<br>Analysis | $\text{cue-evokedAlpha}_n = \beta_{j[n]}^0 + \beta^1 \cdot \text{reward}_{j[n]} \times \beta^2 \cdot \text{caffeine}_n + \beta^3 \cdot \text{session}_n + \epsilon_n$ |
| | $\text{sustained alpha power}_n = \beta_{j[n]}^0 + \beta^1 \cdot \text{caffeine}_n + \beta^3 \cdot \text{session}_n + \epsilon_n$ |

## Author Contributions

B.v.B., M.G.W and M.M.L. designed the study. B.v.B and M.d.J. collected the data. B.v.B and M.d. J. analyzed the data. B.v.B., M.d.J., M.G.W and M.M.L. wrote the manuscript.

